# tDCS modulates effective connectivity during motor command following; a potential therapeutic target for disorders of consciousness

**DOI:** 10.1101/2021.02.09.430392

**Authors:** Davide Aloi, Roya Jalali, Penelope Tilsley, R. Chris Miall, Davinia Fernández-Espejo

## Abstract

Transcranial direct current stimulation (tDCS) is attracting increasing interest as a potential therapeutic route for unresponsive patients with prolonged disorders of consciousness (PDOC). However, research to date has had mixed results. Here, we propose a new direction by directly addressing the mechanisms underlying lack of responsiveness in PDOC, and using these to define our targets and the success of our intervention in the healthy brain first. We report 2 experiments that assess whether tDCS to the primary motor cortex (M1-tDCS; *Experiment 1*) and the cerebellum (cb-tDCS; *Experiment 2*) administered at rest modulate thalamo-cortical coupling in a subsequent command following task typically used to clinically assess awareness. Both experiments use sham- and polarity-controlled, randomised, double-blind, crossover designs. In *Experiment 1*, 22 participants received anodal, cathodal, and sham M1-tDCS sessions while in the MRI scanner. A further 22 participants received the same protocol with cb-tDCS in *Experiment 2*. We use Dynamic Causal Modelling of fMRI to characterise the effects of tDCS on brain activity and dynamics during simple thumb movements in response to command. We found that M1-tDCS increased thalamic excitation and that Cathodal cb-tDCS increased excitatory coupling from thalamus to M1. All these changes were polarity specific. Combined, our experiments demonstrate that tDCS can successfully modulate long range thalamo-cortical dynamics during command following via targeting of cortical regions. This suggests that M1- and cb-tDCS may allow PDOC patients to overcome the motor deficits at the root of their reduced responsiveness, improving their rehabilitation options and quality of life as a result.

## 1. Introduction

Transcranial direct current stimulation (tDCS) is a non-invasive brain stimulation technique that is gaining popularity as a therapeutic option for complex clinical conditions for which no other alternatives are available[1]. Among these, a paradigmatic case is that of prolonged disorders of consciousness (PDOC), such as the vegetative (VS) and the minimally conscious state (MCS). PDOC are characterised by catastrophic disabilities that are in many cases permanent[2], and the small number of therapies available have demonstrated very limited success at improving outcome[3]. In response to this, over the last 5 years the field has seen a sharp rise in tDCS trials on PDOC[4]. These have typically targeted the left dorso-lateral prefrontal cortex (DLPFC), in an attempt to restore some residual level of awareness, but have only had mixed success. While several studies reported the emergence of new behaviours indicative of awareness in subsets of PDOC patients following tDCS (see e.g.[5]), many others have failed to elicit any clinical changes or indeed led to undesired changes[6]. Individual responses to tDCS are well known for their heterogeneity even in healthy populations[7], and we can expect an even higher variability in PDOC, where the specific aetiology and mechanisms of damage result in marked differences in brain atrophy and tissue microstructure across patients. However, in this particular case, we argue that these difficulties are further exacerbated by our limited understanding of how conscious awareness is supported in the brain, which preclude the identification of effective targets for stimulation. Indeed, while we know that consciousness requires sustained rich neural dynamics in fronto-parietal and thalamo-cortical networks[8,9], the specific pattern of activity that would need to be restored in PDOC patients and how this can inform the selection of stimulation targets remains an elusive question.

Here we propose a different approach, wherein we switch the focus from the consciousness disorder itself to the patients’ ability to produce voluntary behavioural responses[10]. In doing so, we target a cognitive process that is much better understood, not only in terms of its neurophysiology but also which specific tDCS modulations can maximise behavioural changes[7]. In addition, recent voices have emphasised the importance of addressing the fundamentals of any tDCS intervention in well-controlled studies in healthy individuals before a clinical application with meaningful effects can be produced and clinically tested[7]. In line with this, we thus focus on characterizing tDCS responses in the healthy brain, while keeping our methods translatable to PDOC patients. Clinical assessments of PDOC use the patient’s ability to follow commands as a proxy measure for their awareness. Crucially, it is well known that a significant number of PDOC patients retain a much greater deal of awareness than can be expected from their clinical diagnosis and are simply unable to demonstrate this with overt purposeful (motor) responses in response to commands[10,11]. We have recently shown that this lack of behavioural responsiveness is associated with specific impairments within the motor system that result in reduced excitatory coupling between the thalamus and the primary motor cortex (M1)[12,13]. On this basis we hypothesise that interventions to enhance the flow of information between the thalamus and motor cortices will provide patients with a renewed level of control over their external behaviour and increase their behavioural responsiveness as a result.

In this study, we use dynamic causal modelling (DCM) of fMRI data to explore whether tDCS can indeed modulate motor thalamo-cortical coupling during simple voluntary responses to command in the healthy brain. We report two separate experiments targeting M1 and the cerebellum respectively. While there is strong evidence that tDCS applied to M1 (henceforth referred to as M1-tDCS) leads to local polarity-specific changes in M1 excitability[14] and BOLD signal[15], little is known about whether it can also influence coupling between other nodes of the motor network. Similarly, there is evidence that cerebellar tDCS (cb-tDCS) is able to modulate cerebellar brain inhibition (CBI)[16], the natural inhibitory tone the cerebellum exerts over M1. Given that the cerebellum is structurally connected to M1 via a thalamic relay, it would follow that the previously reported effects of cb-tDCS on CBI should be mediated by the thalamus. However, no studies have directly investigated how cb-tDCS affects the coupling in this cerebellar-thalamo-M1 axis. Furthermore, no study to date has assessed the effects of either M1- or cb-tDCS on the activity and dynamics of the motor network *during* simple motor command-following. We hypothesised that: (a) anodal M1-tDCS will increase excitation in the motor network and lead to an increased excitatory output from thalamus to M1 during command-following (*Experiment 1*); and (b) cathodal cb-tDCS will reduce inhibition in the thalamus and also result in increased excitation from thalamus to M1 (*Experiment 2*). Previous research has identified a relative structural preservation of M1-striatal-thalamic and dentate-thalamic pathways in PDOC patients[13]. This suggests that both pathways may be viable routes to target the thalamus in this group.

## 2. Material and methods

### 2.1 Participants

Forty-nine right-handed healthy volunteers participated in the study (15 men, 34 women; mean age 25 ± 4 years). We recruited all participants from the University of Birmingham, using the local Research Participation Scheme and advertisements across campus. We pre-screened all participants before recruitment to confirm their eligibility to safely take part in MRI and tDCS experiments. All reported no previous history of neurological and/or psychiatric disorders, no personal or family history of epilepsy, no use of psychoactive drugs, and had normal or corrected vision. Additionally, we instructed them to be well hydrated and well slept, with no alcohol or coffee consumed during the 24 hours prior to the testing session, to be in keeping with brain stimulation safety regulations.[17] The University of Birmingham’s Science, Technology, Engineering and Mathematics Ethical Review Committee approved the study and all participants gave written informed consent prior participation. We compensated participants with £110 or the equivalent in course credits.

*Experiment 1* included 26 participants (8 male, 18 female; mean age 23 ± 4 years), from whom 22 completed all 3 sessions. We further discarded data from one participant due to failure to comply with the task instructions, resulting in a final sample of 21 to be included in the analysis (8 male, 13 female; mean age mean: 23 ± 4 years).

*Experiment 2* included 23 participants (7 male, 16 female; mean age: 27 ± 4 years), from whom 22 completed all 3 sessions. We excluded one further participant due to an acquisition error in one of the sessions that resulted in corrupted files. The final sample consisted of 7 males and 14 females, aged 27 ± 4 years. One participant took part in both Experiments (with a gap of over 7 weeks between them).

### 2.2 General Experimental procedure

Both experiments used sham- and polarity-controlled, randomised, double-blind, crossover designs. All participants completed anodal, cathodal, and sham stimulation sessions, while in the MRI scanner. These were scheduled at least 7 days apart (*Experiment 1*: mean 12 ± <u>10</u>; *Experiment 2:* mean 13 ± <u>7</u>), and in a counterbalanced order. Both the participants and the researchers conducting the data analyses were blind to the polarity in each session.

In their first testing session participants provided informed consent for the <u>study</u> and completed the Edinburgh handedness inventory[18]. Additionally, before each session, we pre-screened participants to confirm MRI and tDCS safety. After completing these steps, we set up the electrodes in a designated room (see below), and took the participants to the MRI scanner, where we completed the setup of the tDCS system and provided the participants with a joystick to record their responses in the fMRI task (see below). We used the MRI Intercom system to communicate with participants during the experiment. Before and after the stimulation, participants performed an fMRI motor command-following task where they were instructed to execute discrete simple thumb movements (abduction-adduction) with their right hand in response to auditory cues (see fMRI paradigm below).

Finally, to test whether our protocol achieved adequate blinding, participants completed a post-tDCS perceptual scale of their perceived sensations and/or discomfort after each session, and indicated whether they thought they received actual stimulation or sham.

### 2.3 Electrical Stimulation

In both experiments we administered tDCS in the MRI scanner using an MR-compatible NeuroConn DC-Stimulator MR (neuroCare Group GmbH, Germany). We used 5×5 cm^2^ electrodes with electro-conductive paste to improve conduction and secured them in place using self-adhesive bandage.

#### Experiment 1

In line with previous studies targeting M1[14], in the anodal sessions we placed the target electrode (anode) centred on the left motor hotspot, as identified by TMS prior to the first MRI session, and oriented approximately at a 45° angle with respect to the midline. We placed the reference electrode (cathode) on the contralateral supraorbital region. We reversed this montage for the cathodal sessions. Half of the sham sessions replicated the anodal montage and the other half the cathodal montage. We used a Magstim BiStim^2^ TMS stimulator paired with Brainsight TMS navigation system (Rogue Research Inc) to identify the motor hotspot in each participant in the first stimulation session, following standard methods[19].

#### Experiment 2

We placed the target electrode on the right cerebellar cortex (3 centimetres lateral to the inion, oriented parallel to the midline) and the return electrode on the right buccinator muscle[20]. The montage was reversed for anodal and cathodal sessions. As above, half of the sham sessions replicated the anodal montage and the other half the cathodal montage.

In both experiments, we used Brainsight to record the coordinates for the target electrode in the first session and used them to locate the electrode position for the subsequent sessions to ensure consistent placement.

During anodal and cathodal sessions, we delivered 20 minutes of stimulation, with 30 seconds of ramp-up and ramp-down periods. During sham, we delivered 30 seconds of stimulation before ramping down to give the sensation of active stimulation, and according to well established protocols to ensure blinding[21]. In *Experiment 1* we stimulated at an intensity of 1mA, as this typically induces tDCS canonical excitatory versus inhibitory effects for anodal and cathodal stimulation respectively[7,14]. In *Experiment 2*, we stimulated at an intensity of 1.85mA as previously recommended[22]. In both studies, we delivered stimulation at rest, without the participant engaging in any motor (or other type of) task, as performing a task during stimulation would not be feasible in PDOC patients themselves.

### 2.4 MRI acquisition

We acquired all data on a Philips Achieva 3T system, with a 32-channel head coil, at the Birmingham University Imaging Centre (BUIC).

#### Experiment 1

fMRI acquisition parameters were as follows: 160 volumes per run, 34 slices, TR = 2000ms, TE = 35ms, matrix size = 80□×□80, voxel size = 3×3×3mm, no gap, and flip angle = 79.1°, SENSE acceleration factor = 2. Additionally, we acquired a high-resolution, T1-weighted MPRAGE image, for anatomical co-registration, with the following parameters: TR = 7.4ms, TE = 3.5ms, matrix size = 256×256mm, voxel size = 1×1×1mm, and flip angle = 7°.

#### Experiment 2

fMRI acquisition parameters were as follows: 119 volumes per run, 46 slices, TR = 2700ms, TE = 35ms, matrix size = 80×80, voxel size = 3×3×3, no gap, flip angle = 79.1°, SENSE acceleration factor = 2. High-resolution, T1-weighted MPRAGE images were also acquired for Experiment 2, with the following parameters: TR = 7.4ms, TE = 3.5ms, matrix size = 256×256, voxel size = 1×1×1, and flip angle = 7°.

In both Experiments we collected other anatomical data as well as resting state fMRI before, during, and after stimulation, but we did not analyse these within the current study, and we will report them in separate papers.

### 2.5 fMRI paradigm

We instructed participants to perform a thumb adduction-abduction movement as fast as they could in response to auditory cues (beeps). The use of a simple task enables both the direct translation of this paradigm to PDOC patients as well as the study of tDCS-induced activation changes independent of modulations of performance. We presented the beeps in blocks cued by the word ‘move’ and interspersed with blocks in which the participant was instructed to rest (cued by the word ‘relax’). Each ‘move’ block included 7 beeps presented at a variable interstimulus interval (range 2-3 seconds), in order to avoid prediction effects. The task included 8 blocks of each type, each with a duration of 20 seconds, and for a total duration of 5 minutes and 20 seconds. We instructed the participants to maintain fixation on a white cross displayed in the centre of a black screen throughout the full duration of the task. This, as well as the instructions at the start of the task (“*Start moving your thumb as quickly as you can every time you hear a beep. Stay still when you hear “relax”. Make sure you keep looking at the fixation cross at all times*”) were presented via a digital system (Barco F35 AS3D, Norway) that projected the image onto a mirror fixed to the head coil at a visual angle of ∼10°. We delivered all auditory cues with an MR-compatible high-quality digital sound system incorporating noise-attenuated headphones (Avotec Silent Scan®). During ‘move’ blocks, we recorded thumb movements with an MRI compatible joystick (FORP-932, Current designs INC., PA USA), using 1200 Hz sampling frequency of x and y positions. To facilitate use of the joystick inside the MRI bore, the device was connected to the interface in the control room through an optical cable. For each session, we stabilised the joystick on the participant’s torso and stabilised their right thumb using tape. To ensure accurate recordings, we calibrated the joystick before starting the experiment in each session. We used MATLAB 2015b on a Windows 7 computer to deliver all task stimulus and record motion tracking. See Fig. 1.

**Fig. 1.**
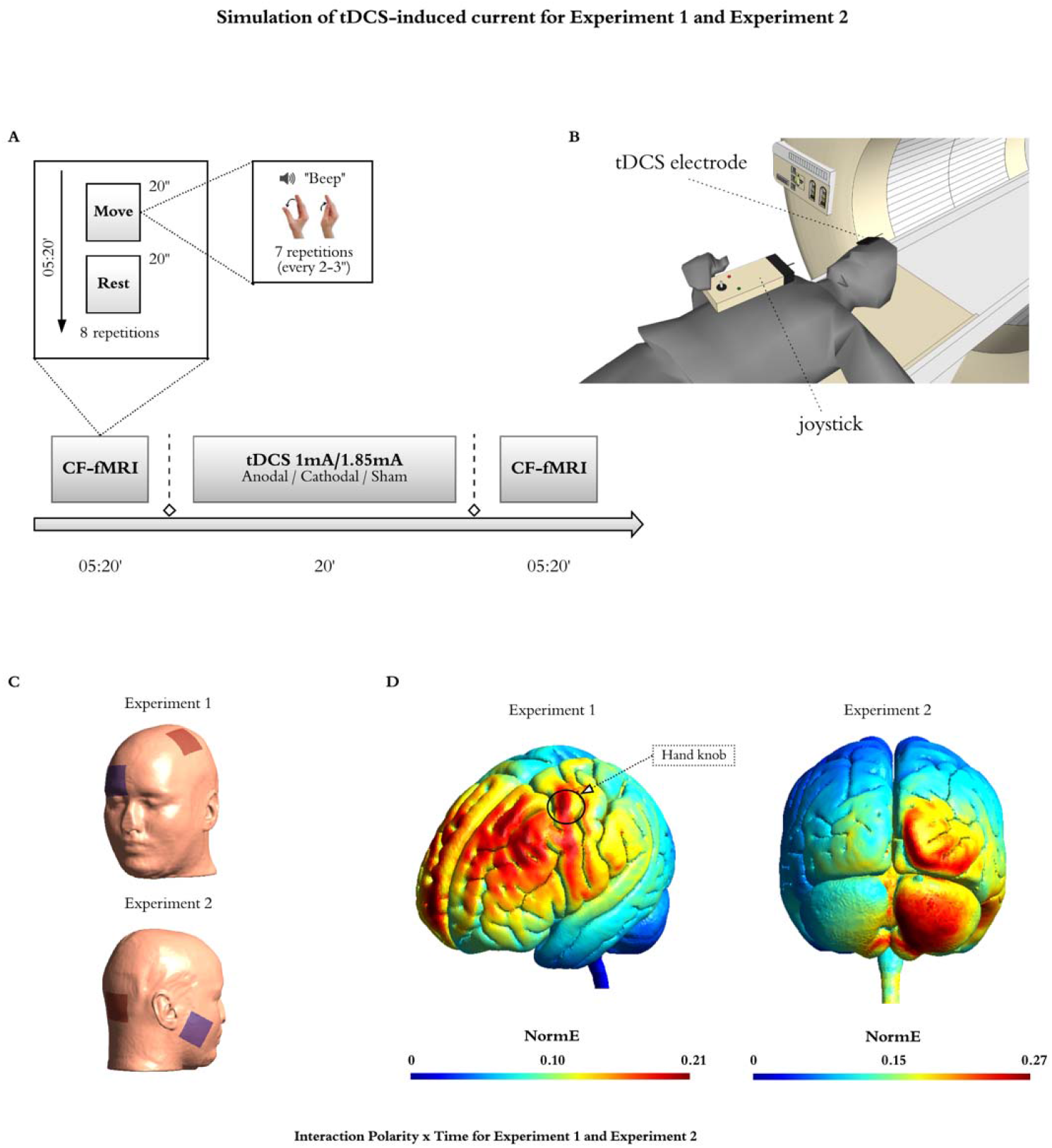
Experimental Design and tDCS montages. Experimental design. (**A**) Participants performed a simple behavioural command following task in the MRI scanner (CF-fMRI) before and after receiving 20 minutes of tDCS, whereby they move their right thumb in response to auditory cues (beeps). The task alternated 8 blocks of movement interspersed with rest blocks (all blocks were 20 seconds long for a total of 5 minutes 20 seconds). The beginning of each block was cued by the auditory words ‘move’ (movement blocks) or ‘relax’ (rest blocks). In each ‘move’ block the participants were instructed to perform 7 discrete thumb adduction-abduction movements as fast as they could in response to beeps that appeared at intervals ranging from 2-3 seconds, and while keeping their gaze fixated on a fixation crossed displayed in the centre of a black screen. Their movements were recorded with an MRI compatible joystick, using 1200 Hz sampling frequency of x and y positions (**B**). All participants received anodal, cathodal, and sham stimulation sessions in a counterbalanced order at least 7 days apart. In Experiment 1, we used a montage that targeted the left primary motor cortex (M1) with the reference electrode over the contralateral supraorbital region, and delivered our stimulation at 1mA (**C, top inset**). We used TMS to identify the best placement (motor hotspot) of the active electrode in each participant. In Experiment 2, our montage targeted the right cerebellar cortex, with a reference electrode over the right buccinator muscle, and delivered our stimulation at 1.85mA (**C, bottom inset**). (**D)** Computational model showing the electric field distributions in Experiment 1 (left) and Experiment 2 (right), as calculated with SimNIBS3.2.2 on the MNI standard head model. For the purpose of this simulation, in Experiment 1, we placed the active electrode on C3 to approximate the location of the motor hotspot in our participants (marked as hand knob in the figure), and the passive electrode on Fp2. In Experiment 2, we placed the active electrode on I2 and the passive electrode over the right buccinator muscle. Note that this model does not consider individual differences in the position of the electrodes or the different tissue compartments across individual participants and therefore it should be interpreted as an estimate of the canonical field distribution to be expected with our montages.

### 2.6 fMRI preprocessing and GLM analysis

We used SPM12 on MATLAB 2015b (www.fil.ion.ucl.ac.uk/spm) for the preprocessing and analysis of both fMRI datasets.

Each dataset was analysed independently but following the same pipeline, as described here. We first followed a standard spatial pre-processing, including realignment, co-registration between the structural and functional data sets, spatial normalization, and smoothing with an 8mm fwhm Gaussian kernel). Additionally, in order to remove potential undesirable effects of physiological noise or participant’s motion in the activation maps, we performed single-subject independent component analysis (ICA)[23] and then applied FMRIB’s ICA-based X-noiseifier (FIX)[24,25] to identify artefactual components and remove them from our fMRI data. We first classified manually all components from a subset of datasets (18 in *Experiment 1* and 23 in *Experiment 2*), ensuring an even coverage of all possible combinations of sessions, times, and polarities. Then, we used these manual labels to train a classifier for each of the studies that we then applied to the remaining datasets in that study. In order to test the accuracy of the automatic component classification, two of the authors (D.F-E for *Experiment 1* and D.A. for *Experiment 2*) independently classified a number of components in the training set (8 datasets for *Experiment 1* and 10 datasets for *Experiment 2*) and cross-checked their manual classification against the automatic classifications performed by FIX. There was a 100% match for ‘bad’ components between the manual and automatic classification lists.

We performed single-participant fixed-effect analyses using a general linear model in which we modelled each scan to belong to the motor execution (i.e. blocks of thumb movements) or the rest condition. The model also included the realignment factors as effects of non-interest to account for residual motion-related variance. We used high-pass filtering with a cut-off period of 80 seconds to remove slow-signal drifts from the time series. We then set linear contrasts to obtain estimates of the effects of interest for each subject, polarity, and time. Finally, in order to test the effects of tDCS on brain activation, we performed a second level full factorial analysis with polarity (anodal, cathodal, and sham) and time (before and after tDCS) as factors (total number of sessions = 126 for *Experiment 1* and 126 for *Experiment 2*). When the interaction was significant, we also performed the corresponding pairwise interactions to study the direction of the effects. We report statistically significant voxels as being those that survive an uncorrected p<.0001 at the voxel level, on the following regions of interest: left supplementary motor area (SMA), left precentral gyrus, left thalamus, and right cerebellar lobes IV-V and VIII[26], using WFU PickAtlas. We did not include spurious activation, defined as a contrast returning a single significant voxel. We obtained these regions of interest from the Automated Anatomical Labeling atlas[27]. In *Experiment 1*, we had to exclude one participant from the ANOVA due to an acquisition error in one of the sessions that resulted in the most superior slices of the brain being cropped (losing a small section of M1). Note however that this issue did not affect the VOI analyses for the DCM (see section below) and therefore this participant was included in the DCM analyses. See full analysis pipeline in Fig. 2.

**Fig. 2.**
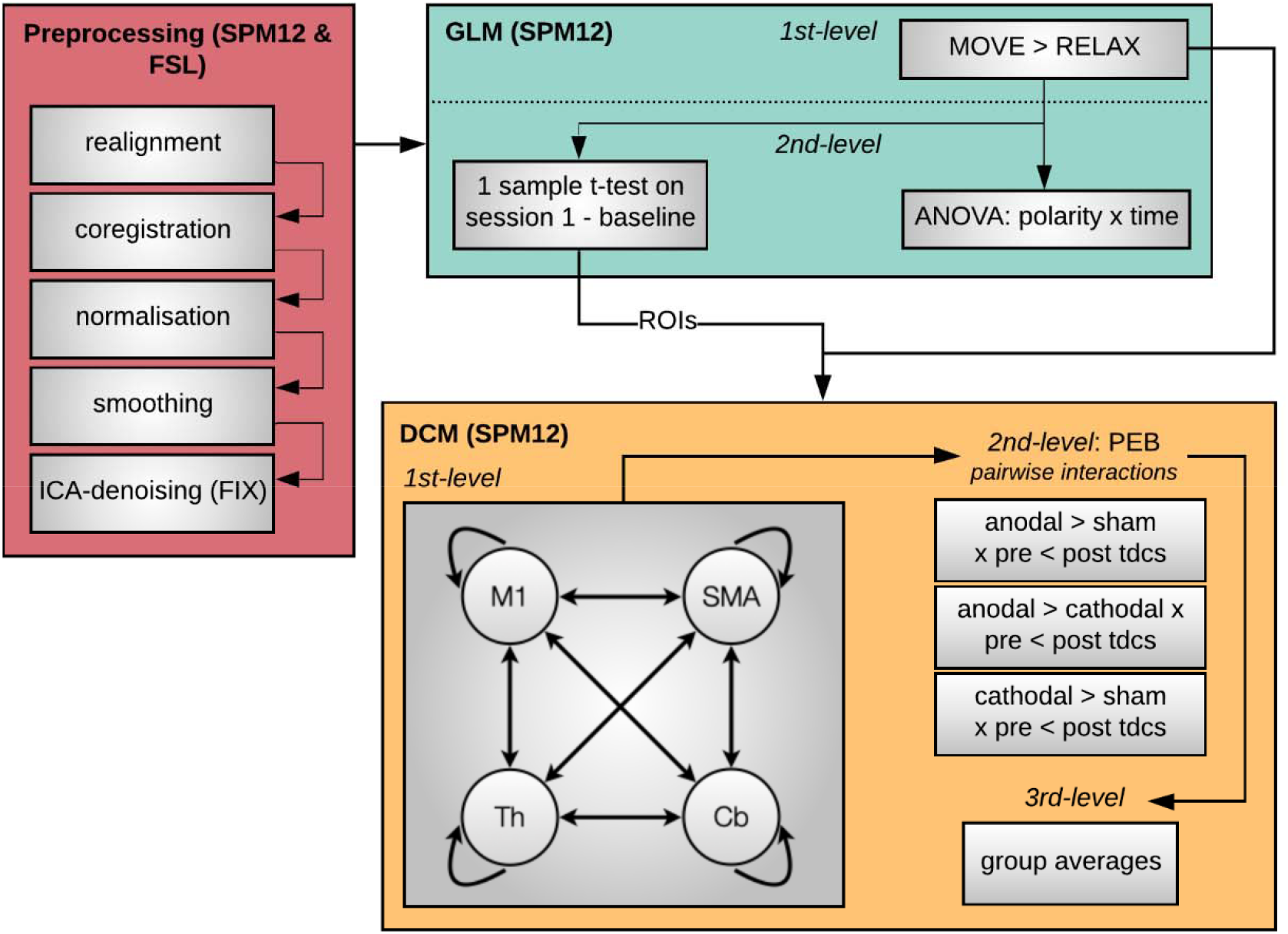
Analysis pipeline. Analysis pipeline. We followed a standard pre-processing protocol (red panel), followed by fixed-effect general linear model analysis to model the effect of thumb movements in each individual participant (1st-level, green panel). We then conducted a second level full factorial analysis to test the effects of tDCS on brain activation (green panel). In addition, we performed a second-level one-sample t-test on the pre-stimulation run acquired in the first chronological session for each participant to characterise the canonical activation in the task, and define coordinates for the subsequent dynamic causal modelling (DCM) analyses. Finally, we used DCM to assess the effects of tDCS on the causal dynamics within our network of interest (yellow panel). We first built and estimated a fully connected model including left M1, left SMA, left thalamus, and right cerebellum in each participant. Then we applied Parametric Empirical Bayes (PEB) to model each of the three pairwise interactions between polarity and time (i.e., interaction between pre-/post-tDCS and either anodal/cathodal, cathodal/sham, anodal/sham) in each participant (2nd-level, yellow panel). Finally, we created a 3rd-level PEB for each pairwise interaction modelling the average effect across participants. Note that we conducted data analysis for each Experiment individually but following the same protocol, as described above.

### 2.7 DCM analysis

#### Region selection and timeseries extraction

DCM is a framework for Bayesian modelling of brain dynamics, which allows the inference of hidden (unobserved) neuronal states from measured brain activity [28]. First, to obtain the canonical pattern of activity on our task for the group in each experiment, we performed second-level one-sample t-tests on the individual contrasts corresponding to the pre-stimulation run acquired in the first chronological session for each participant (Fig. 3). In the resulting map, we identified the group peak of activation for the clusters corresponding to the left M1, SMA, left thalamus, and right cerebellum at an uncorrected p<0.001 (in bold in Table 1). This group-derived coordinates then served as a starting point for searching a nearby local maximum in each individual run. Each of these run-specific local maxima was constrained to be a maximum of 15mm away from the group level peak for the left M1, SMA, and right cerebellum ROIs and a maximum of 9mm away for the left thalamus ROI, and had to exceed a liberal statistical threshold of p<0.05[28]. The differences in the allowed distance from the group peak accommodated for differences in size of the anatomical boundaries of each region. As recently recommended, when this threshold failed to produce a peak for that region, we iteratively reduced the threshold in 0.05 increments until reaching 0.25. When no peak could be found even at this threshold, we used the original group derived coordinates, as typically done[29]. Note that we only used these liberal thresholds for the identification of coordinates to extract our timeseries (feature selection) but not for any statistical analyses. Having identified individual peak coordinates for each run, we extracted timeseries from 4mm radius spherical volumes of interest centred on them.

**Table 1.** Canonical activation during command-following.

|  | Region | Cluster P | Cluster size | Peak P | T | MNI coordinates |
| --- | --- | --- | --- | --- | --- | --- |
|  |  | *FWE-corrected* | *in mm3* | *uncorrected* |  | *[x;y;z]* |
| Experiment 1 | M1 | <0.001 | 6264 | <0.001 | 6.770 | **[-33;-13;62]** |
|  |  |  |  | <0.001 | 6.767 | [-33;-19;56] |
|  |  | 0.966 | 81 | <0.001 | 4.751 | [-15;-4;68] |
|  |  | 0.903 | 162 | <0.001 | 4.704 | [-57;5;29] |
|  |  | 0.927 | 135 | <0.001 | 4.324 | [-54;5;14] |
|  | SMA | <0.001 | 6426 | <0.001 | 8.703 | **[0;-7;65]** |
|  | Thalamus | 0.260 | 864 | <0.001 | 6.330 | **[-12;-22;5]** |
|  | Cerebellum | 0.022 | 2187 | <0.001 | 7.137 | **[15;-55;-22]** |
| Experiment 2 | M1 | 0.326 | 702 | <0.001 | 5.066 | **[-24;-19;74]** |
|  |  | 0.983 | 81 | <0.001 | 4.565 | [-54;5;14] |
|  |  | 0.983 | 81 | <0.001 | 4.251 | [-15;-4;68] |
|  |  | 0.195 | 918 | <0.001 | 4.107 | [-42;-19;56] |
|  |  |  |  | <0.001 | 4.085 | [-36;-28;65] |
|  |  |  |  | <0.001 | 3.957 | [-33;-25;53] |
|  |  | 0.983 | 81 | 0.001 | 3.795 | [-39;-7;47] |
|  |  | 0.997 | 27 | 0.001 | 3.606 | [-33;-22;47] |
|  | SMA | <0.001 | 5481 | <0.001 | 7.087 | **[-3;-4;65]** |
|  |  |  |  | <0.001 | 6.178 | [-12;-4;74] |
|  |  |  |  | <0.001 | 5.345 | [-9;-1;53] |
|  | Thalamus | 0.505 | 513 | <0.001 | 4.352 | **[-18;-16;14]** |
|  |  |  |  | <0.001 | 4.153 | [-9;-22;5] |
|  | Cerebellum | 0.003 | 3024 | <0.001 | 8.799 | **[12;-55;-25]** |
|  |  |  |  | <0.001 | 7.671 | [18;-49;-19] |
|  |  |  |  | <0.001 | 5.159 | [24;-43;-31] |
|  |  | 0.001 | 3915 | <0.001 | 7.285 | [24;-58;-46] |
|  |  |  |  | <0.001 | 7.255 | [12;-73;-46] |
|  |  |  |  | <0.001 | 6.467 | [6;-67;-31] |

Results from the random effect group analyses on the brain activation during thumb movements to command in the baseline run for the first session. Results survived a threshold of uncorrected $p < 0.001$ . We highlight in bold the coordinates that we subsequently used as a starting point to search for individual coordinates to extract time series for the DCM. Abbreviations: FWE, family wise error; MNI, Montreal Neurological Institute; M1, primary motor cortex; SMA, supplementary motor area.

**Fig. 3.**
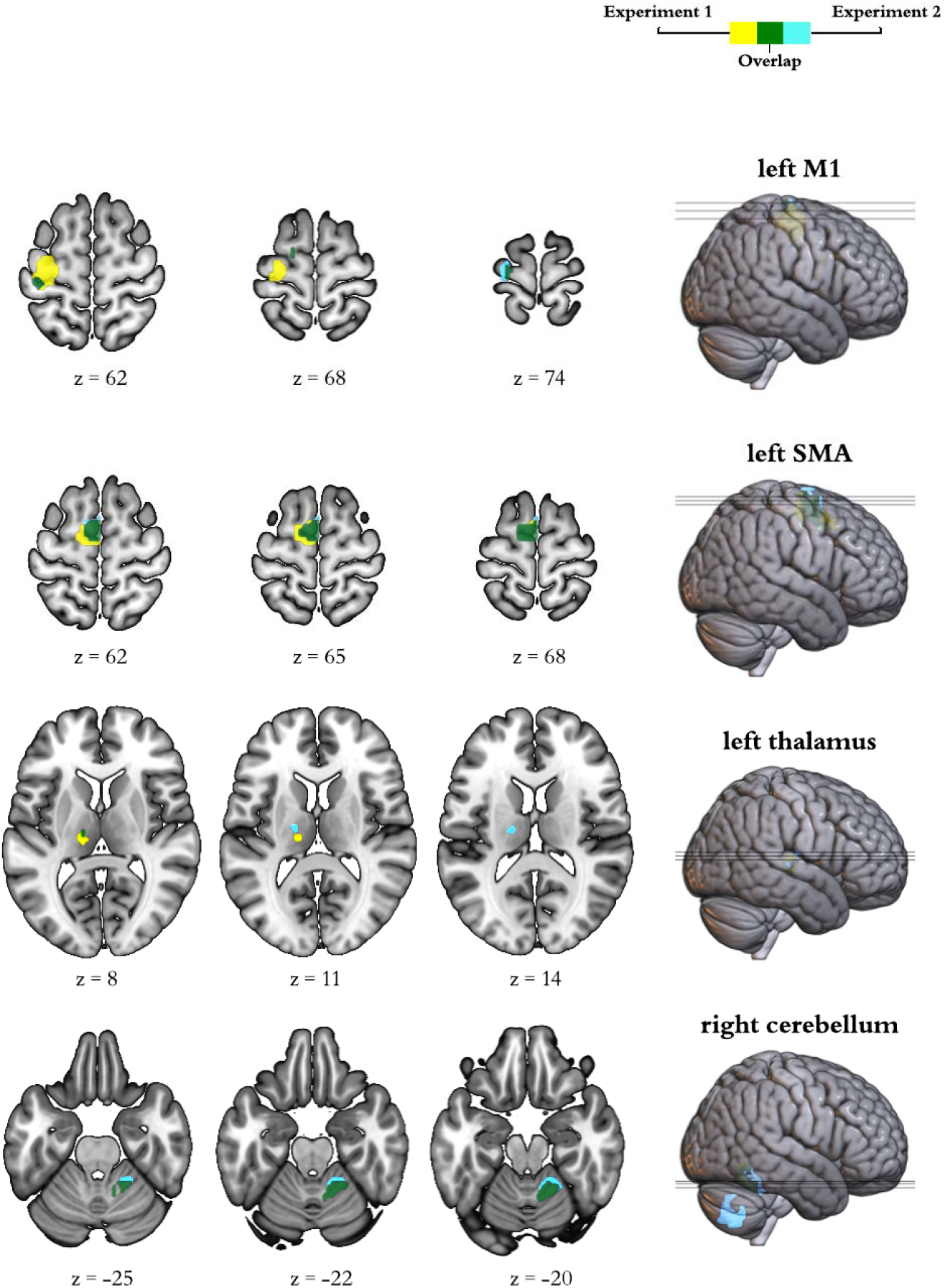
Activation at Baseline. Brain activation during command following in the pre-stimulation run corresponding to the first session for each participant. The insets display group general linear model differences between ‘move’ and ‘rest’ blocks in Experiment 1 (yellow) and Experiment 2 (light blue). The overlap across experiments appears in green. For display purposes, activation maps are shown at an uncorrected p<0.001 and rendered on a standard template (152 template in MRIcroGL). z indicates the Montreal Neurological Institute z coordinate.

#### 2.7.1 Individual level DCM specification and definition of model space

With the above extracted timeseries, we specified individual dynamic causal models using the deterministic model for fMRI, one-state per region, bilinear modulatory effects, and mean-centred inputs. We started with a 4-node fully connected model in which all self- and between region connections were switched on. The effect of thumb movements entered the model as modulatory input on the self-connection of each region, as this is recommended to improve both parameter identifiability and biological interpretability [28]. In addition to the intrinsic connections and modulatory inputs above, DCM requires the specification of driving inputs, which briefly ‘ping’ specific regions in the network at the onset of each block. In order to determine the best set of inputs for our data, we first created DCMs that included driving inputs to all 4 regions in our model and applied Parametric Empirical Bayes (PEB) to prune any parameters that were not contributing to the model evidence. Briefly, PEB is a hierarchical Bayesian framework for group-level modelling of effective connectivity, that allows the evaluation of both group effects and between-subject variability over DCM parameters (see [29] for a full description). For this step, we created a second-level PEB modelling the commonalities across all 6 sessions for each participant. These were then fed to a third-level PEB that modelled the commonalities across the group. In addition to the constant encoding the group mean, we included sex, age, and the score in the Edinburgh Handedness Inventory as nuisance regressors (all mean-centered). Finally, we used Bayesian Inference to invert the model for each subject and estimate the parameters that maximise explanation of data while minimising complexity. For this, we used Bayesian Model Reduction (BMR) to search over the reduced models followed by Bayesian Model Average (BMA) to calculate the average connectivity parameters[29]. We used a 95% posterior probability threshold for free-energy (i.e., comparing the evidence for all models where a particular connection / input is on, versus those where it is off). This step indicated strong support (>99% posterior probability) for including driving inputs to cortical regions (M1, and SMA) only (see results for full details) and therefore we re-defined DCMs for all of our participants using these parameters. Our final model therefore included all self- and between-region connections, modulatory inputs to each self-connection, and driving inputs to M1 and SMA.

#### 2.7.2 PEB ANOVAs

To test the effects of tDCS on the model parameters (connections and task modulations), we first created 3 second-level PEB models in each participant, which encoded the following pair-wise interactions: (1) greater increases after anodal stimulation as compared to sham (pre-tDCS < post-tDCS x anodal > sham sessions) and (2) greater increases after anodal stimulation as compared to cathodal (pre-tDCS < post-tDCS x anodal > cathodal), and (3) greater increases after cathodal stimulation as compared to sham (pre-tDCS < post-tDCS x cathodal > sham). Note that these contrasts also encode the opposite effects: e.g., PEB 1 can also be interpreted as greater decreases in sham as compared to anodal (pre-tDCS > post-tDCS x anodal < sham). Each subject specific PEB model was then entered into one of 3 third-level PEBs that encoded the commonalities across the group (mean) for each pairwise interaction, as well as sex, age, and handedness score.

We then used BMR and BMA to prune connections that do not contribute to the model evidence and estimate the parameters across all models for each of the connections that remain switched on. We thresholded our BMA results at a posterior probability > 95% (which is equivalent to a Bayes factor of 3) [29].

### 2.8 Motion tracking

We performed motion data analysis using a custom script on MATLAB 2017b. First, we calculated the Euclidean distance of the x-y position and applied a low-pass 15Hz filter to the data. We then identified the onset and end of the movement by looking at abrupt changes in the signal, using the matlab function *findchangepts*, which, given a vector x with N elements (in our case containing motion tracking data) returns the index at which the mean of x changes most significantly. We used the first and last change detected by *findchangepts* to determine when each movement started and ended. We excluded movements where no changes were detected, which could be due to participants not responding to the task or to the joystick not recording data. In *Experiment 1*, this resulted in the removal of 5 datasets from the motion tracking analysis, due to the joystick malfunctioning during recording in at least one of three sessions. Lastly, we calculated velocity and acceleration at each timepoint between the beginning and end of each movement and obtained the mean velocity and peak acceleration for the trial. Additionally, we calculated reaction time defined as the time occurring between the auditory stimulus (beep) and the onset of the movement. Finally, we averaged these values across each run and computed a 2 (pre-vs post-tDCS) x3 (polarity) repeated measures ANOVA to check for any effect of tDCS on behaviour.

### 2.9 Blinding

In order to assess whether our blinding protocol was successful, in each Experiment, we used McNemar’s test to assess whether the number of correct judgements across the group about whether they had received tDCS or not was different between real stimulation and sham stimulation sessions.

## 3. Results

### 3.1 Experiment 1 - Effects of M1-tDCS on brain activation and dynamics

See the canonical task activation at baseline in Table 1 and Fig. 3.

Our factorial analysis on the individual activation maps revealed a significant interaction between polarity (anodal, cathodal, and sham) and time (pre-, post-tDCS) on the left thalamus only (uncorrected p<0.001; see Table S1 and Fig. 4). Subsequent pairwise interactions revealed that both anodal and cathodal stimulation increased activity in this area as compared to sham, with no significant differences between polarities. (See Supplementary Table S1 and Figure S1 for the positive effect of the task across all sessions in this ANOVA).

**Fig 4.**
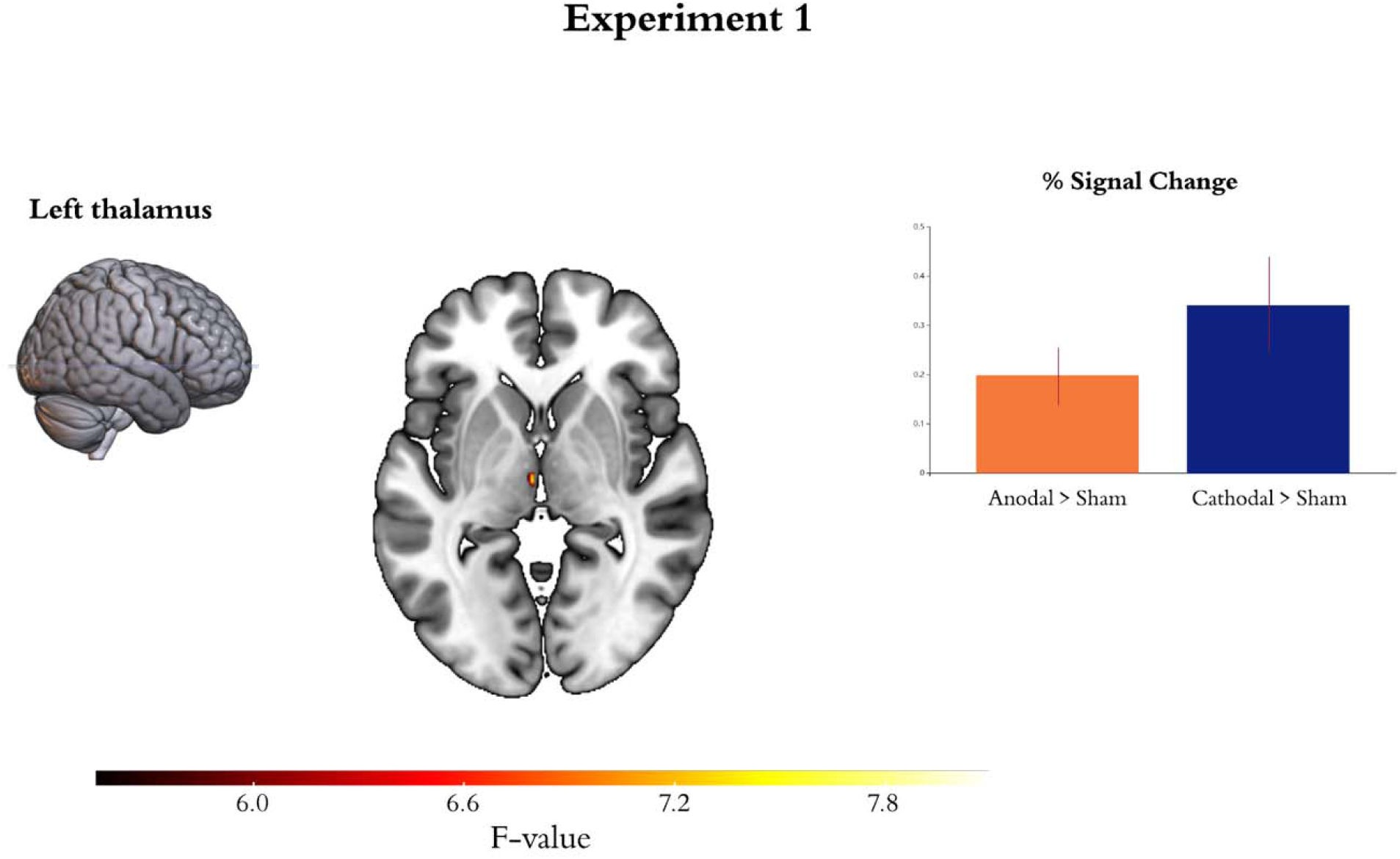
Effects of tDCS on brain activation during command following for Experiments 1. The brain inset display group general linear model (GLM) interactions between polarity (anodal, cathodal, sham) and time (pre-, post-tDCS) in the individual contrasts modelling brain activation during command following. For display purposes, activation maps are shown at an uncorrected p<0.005 and rendered on a standard template (152 template in MRIcroGL). The colour bar represents the F value for the interaction in the GLM. *z* indicates the Montreal Neurological Institute z coordinate. Bar plots show the estimated effect size and 90% confidence intervals at the peak voxel for each pairwise contrast: greater activation after anodal stimulation as compared to sham (orange), and greater activation after cathodal stimulation as compared to sham (blue).

Our DCM analyses revealed that anodal stimulation of M1 reduced self-inhibition in the thalamus and led to a more inhibitory output from cerebellum to M1, compared to both sham and cathodal stimulation. Additionally, as compared to sham, anodal stimulation increased inhibition in all outputs from M1 to the rest of the network but reduced inhibition from cerebellum to thalamus, as well as in SMA and cerebellar self-connections. These changes were however not polarity specific. In turn, cathodal stimulation increased excitation from thalamus to SMA, both as compared to sham and to anodal stimulation. Additionally, as compared to sham, cathodal stimulation led to an increase in inhibition from both M1 and cerebellum to SMA, an increase in excitation from thalamus to M1, and a reduction in self-inhibition in SMA. In terms of task modulations, cathodal M1 stimulation increased the modulatory input from the task on M1 (i.e., increased M1 self-inhibition) both as compared to anodal stimulation and sham, and decreased the modulatory input from the task on SMA as compared to anodal stimulation (see Fig. 5).

**Fig 5.**
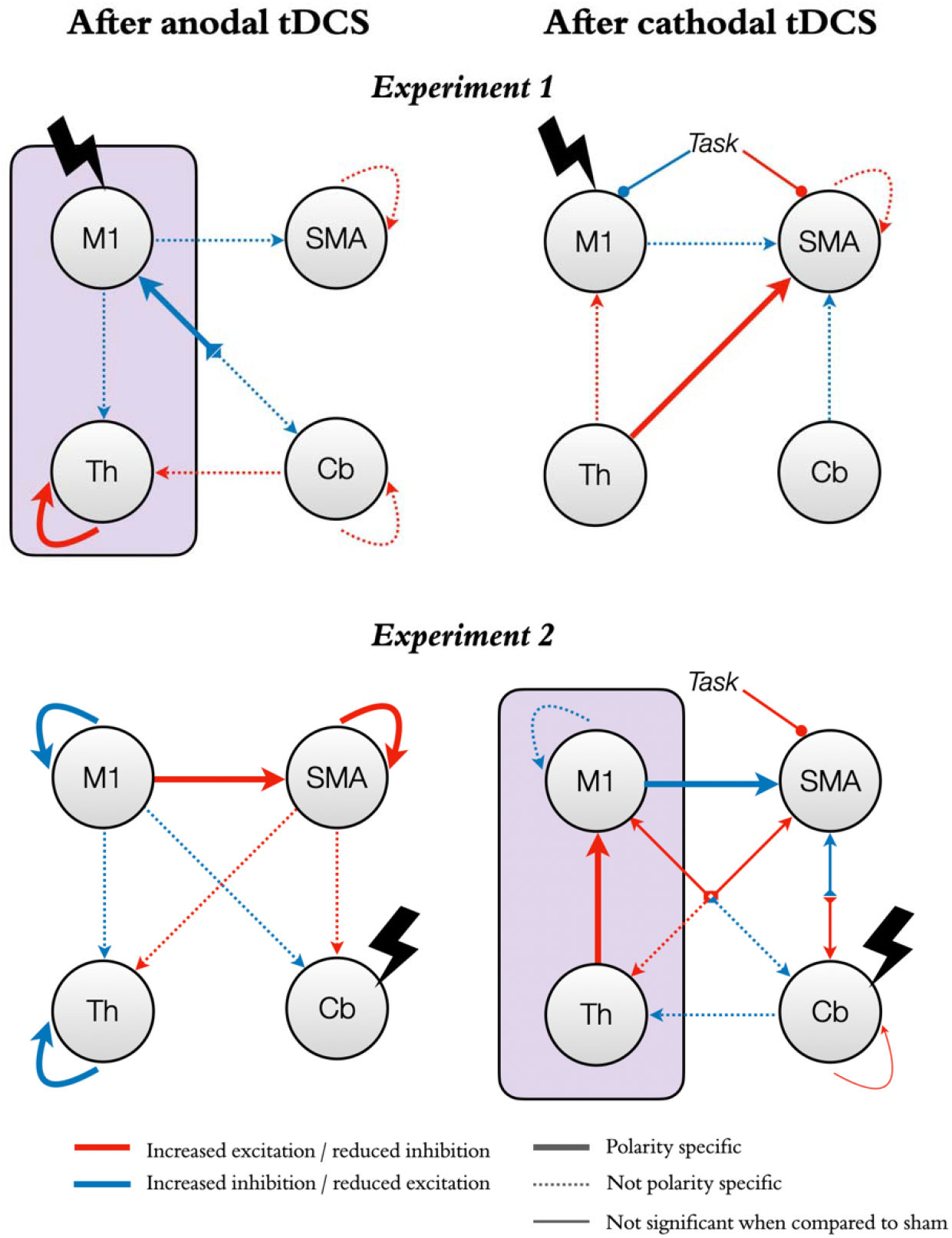
Effects of tDCS on functional neural dynamics with M1 or the cerebellum as targets. The figure shows the effects of tDCS on functional neural dynamics for the two experiments (experiment 1, top panels; experiment 2, bottom panels). The left and right panels represent changes after anodal and cathodal stimulation respectively. Red arrows indicate changes in the direction of increased excitation (or reduced inhibition). Blue arrows indicate changes in the direction of increased inhibition (or reduced excitation). Note that self-connections are always inhibitory and thus red indicates a reduction in inhibition rather than an excitatory role per-se. Similarly, modulatory inputs from our command following task on each region increase (blue) or decrease (red) the region’s inhibitory tone. Thick lines represent changes that are significant both as compared to the opposite polarity and to sham. Thin lines represent changes that are only significant as compared to the opposite polarity. Dashed lines represent changes that are only significant as compared to sham (not polarity specific). The purple boxes highlight our hypotheses for the M1-thalamus axis: anodal M1-tDCS (top left panel) reduced self-inhibition in the thalamus while cathodal cb-tDCS (bottom right panel) increased excitation from thalamus to M1, both in a polarity specific manner.

### 3.2 Experiment 2 - Effects of cb-tDCS on brain activation and causal dynamics

In terms of brain activity during command following, our factorial analysis revealed no significant interactions between polarity and time (pre-vs post-tDCS) in any of the ROIs. See Supplementary Table S1 and Figure S1 for the positive effect of the task across all sessions in this ANOVA.

In terms of effective connectivity, as predicted, cathodal stimulation led to increased excitation from thalamus to M1 both as compared to sham and anodal stimulation. In addition, it increased M1 self-inhibition as compared to sham but to a lesser extent than anodal stimulation. Finally, it increased inhibition from M1 to SMA both as compared to sham and anodal stimulation (Fig. 5). When compared directly with anodal stimulation, cathodal cb-tDCS also decreased cerebellar self-inhibition and increased excitation from thalamus to SMA. Additionally, cathodal stimulation increased inhibition from M1 to cerebellum and from cerebellum to thalamus, and increased excitation from SMA to both thalamus and cerebellum, and from cerebellum to M1, as compared to sham. However, none of these changes were significant when compared with anodal stimulation. Finally, cathodal cb-tDCS decreased the effect of the task on SMA both as compared to sham and anodal stimulation. In contrast, anodal stimulation, when compared to sham and cathodal stimulation, led to increased self-inhibition in M1 and thalamus, reduced self-inhibition in SMA, as well as increased excitation from M1 to SMA. Additionally, anodal stimulation increased excitation from SMA to thalamus and cerebellum, and increased inhibition from M1 to cerebellum when compared to sham, but these changes were not polarity specific (i.e., did not reach statistical significance in the comparison between anodal and cathodal stimulation).

### 3.3 Experiments 1 and 2 - Effects of tDCS on behaviour

As expected, we did not find any interactions (polarity x time) for any of the metrics considered in Experiment 1 nor 2 (i.e., reaction time, mean velocity, and peak acceleration). In Experiment 2 only, we found a small main effect of time on average reaction times, which was 0.02 seconds (20 ms) faster in the second run as compared to baseline (pre-: 0.30s ± 0.04; post-tDCS: 0.28s ± 0.04; p<0.001 uncorrected, α_p_^2^0.5). See Supplementary Table S2 for full statistical information for all main effects, interactions, and post hoc tests.

### 3.4 Experiments 1 and 2 - Blinding

We found no significant differences in the number of times that sham and active stimulation sessions were perceived as real in either experiment, suggesting that participants’ experiences did not differ between active and sham stimulation sessions and blinding was successful (see Supplementary Table S3).

## 4. Discussion

Efforts to use tDCS as a therapeutic intervention in PDOC have had mixed success to date. While some studies showed very promising clinical improvements, many others failed to show any effects even after repeated sessions[30]. The field is thus unable to reach a consensus about whether tDCS would or would not be a feasible therapeutic avenue for this patient group as a result. Most research to date has focused on targeting the left frontal cortex, in an attempt to engage non-specific networks involved in arousal and awareness. Here, we propose a new therapeutic direction that directly addresses the neural mechanisms that support measurable changes in behavioural responses after tDCS at the level of functional thalamo-cortical coupling within the motor network[12].

Our results provide the first evidence that tDCS over motor areas can distally modulate brain activity and causal dynamics in thalamo-cortico-cerebellar loops (beyond the immediate target area of stimulation) *during* behavioural command following, even when the stimulation is delivered at rest. In Experiment 1, anodal stimulation over M1 increased task-induced activation the thalamus. Our DCM analyses revealed that this is likely explained by reduced thalamic self-inhibition. In Experiment 2, cathodal cerebellar stimulation did not lead to changes in task-induced activity but instead led to increased excitatory influence from thalamus to M1. Taken together, these experiments demonstrate that it is possible to influence thalamo-cortical coupling indirectly via targeting *surface* (easily accessible) regions in the motor network. More importantly, they suggest that this could be a viable route to elicit clinically relevant changes in PDOC. Indeed, we designed our command-following task to emulate the approach that is routinely used in clinical settings to assess awareness after severe brain injury; namely asking the patient to perform a discrete movement in response to a verbal command[31]. This resulted in a task that was insensitive to potential tDCS modulations of behaviour in healthy participants but allowed us to study the neural effects of tDCS independently of performance, permitting us to draw more direct comparisons to the PDOC population. Specifically, our task deviated from those typically used in the motor learning literature (e.g., [32,33]) in three crucial points: the use of a very small number of trials (approximately 80-90% less), variable cue intervals to avoid prediction effects, and no feedback to participants. Further, we delivered stimulation at rest to increase the translatability of our results to unresponsive PDOC patients. It is important to highlight that the aim here was not to improve motor control in the healthy brain. Instead, we built upon convincing evidence that the thalamus is greatly inhibited in PDOC due to both structural and functional damage[34–36], resulting in less cortical excitation[36]. Our focus thus lay on compensating for this thalamic over-inhibition instead of enhancing normal function. We have previously shown that increased thalamic activity and excitation, as well as increased excitatory thalamus-M1 coupling facilitates the production of motor responses to command in tasks like the one we used here[12]. We now show that anodal tDCS over M1 and cathodal tDCS over the cerebellum can each modulate these dynamics, albeit in different ways, and we propose that they may allow PDOC patients to overcome motor control deficits at the root of their diminished behavioural responsiveness[12,13]. This in turn would allow more patients to demonstrate their true level of awareness, especially in those affected by so called cognitive motor dissociations[10]. Alongside ensuring that each patient receives an appropriate diagnosis, this increased responsiveness can also have important implications for prognosis by facilitating patients’ engagement with rehabilitation[37]. Moreover, regaining some level of control over their thumb would facilitate the use of assistive devices (including those for communication), which could have an enormous impact on their quality of life. Indeed, to further increase the clinical relevance of our study, we focused on thumb movements, as they are affected by spasticity in fewer PDOC patients and with less severity as compared to other fingers[38].

Importantly, our results suggest two potential routes to target the thalamo-M1 axis, providing some flexibility to adapt the tDCS montage to the specific pattern of injuries present in each individual patient. Crucially, while many PDOC patients present localised structural damage to the white matter fibres connecting thalamus and M1[12,13], this damage is partial instead of a complete deafferentation[13]. This suggests the remaining pathways may be amenable to therapeutic intervention. In contrast, the white matter pathways connecting the cerebellum with the thalamus appear relatively preserved[13], suggesting that this may be a feasible route into the thalamus in the majority of PDOC patients. We have previously argued that the relative preservation of this pathway, in the context of damage to the thalamus and the white matter fibres connecting thalamus to M1, may be contributing to excessive thalamic inhibition[13]. As discussed above, our current results show that cathodal cb-tDCS may be able to successfully counteract this. It is important to acknowledge here that, while both anodal M1 and cathodal cb-tDCS successfully modulated thalamic activity, there were differences in their respective effects over M1 activity and the thalamo-M1 dynamics. Furthermore, cathodal M1-tDCS also led to changes in thalamo-M1 coupling in the desirable direction (increased coupling), alongside increases in thalamic activity. This adds further support to the now well accepted notion that the two polarities do not always result in opposing effects [7]. We include below discussion of potential compensatory mechanisms that may explain these effects, but we cannot rule out that cathodal M1-tDCS may also have therapeutic effects in some PDOC patients. We also note the possibility of simultaneous anodal-M1 and cathodal-cerebellar stimulation, although we have not tested this montage. In any case, further studies in PDOC patients themselves are required to test which of these modulations has greater therapeutic effect and for which specific patients. More broadly, while our results provide a robust proof-of-principle for the use of motor tDCS in PDOC, the specific dose, duration, and number of sessions required to induce reliable neural and behavioural changes in PDOC patients needs to be established. Further, the effects of tDCS are highly variable across individuals [39] and this heterogeneity can only be expected to be greater in PDOC patients, due to individual differences in brain damage affecting thalamo-cortical regions and their structural connectivity [13,35,40]. We report here group effects and thus our results cannot be interpreted in terms of M1 or cerebellar tDCS resulting in less (or more) individual variability as compared to other available interventions (e.g., DLFPC). Indeed, an exploration of individual tDCS differences and their relationship to individual brain structure and white matter connectivity is beyond the scope of the current study but remains a crucial area of further investigation. By focusing on specific circuits that have a mechanistic role in PDOC, we believe our study provides a framework to study individual effects in a robust way.

To our knowledge, only 3 studies have targeted motor areas with tDCS in PDOC[41–43], in sharp contrast with the many others that have focused on the DLPFC, and currently represents the main direction in the field. These 3 motor studies included a combined total of 40 patients (14 VS and 26 MCS). Their small sample sizes, key differences in specific montages and stimulation parameters, alongside the focus on behaviour instead of neural markers, preclude us from drawing direct comparisons with our study. In addition, while we are satisfied that we were able to identify the optimal location of the electrodes on the scalp to target the desired regions in our study, this is a much more challenging task in patients with severe brain injury, where large macrostructural changes will affect the relative position of the brain structures of interest in respect to the scalp. Nevertheless, PDOC studies provided preliminary evidence that M1 and cerebellar tDCS are well tolerated in this patient group and can indeed lead to specific improvements in motor responsiveness in a subset of patients (as indexed by increases in the motor and auditory CRS-R subscales).

Beyond the immediate implications for the rehabilitation of PDOC patients, our results speak for the ability of tDCS to influence long-range dynamics in the motor network during movement execution. The field of non-invasive brain stimulation has recently been tainted by a certain level of scepticism towards the effectiveness of tDCS, with some questioning whether it is indeed capable of modulating brain function at all[39]. The increasing number of well controlled imaging and electrophysiological studies has provided reassurance that tDCS can indeed modulate cortical regions under the electrodes. In the specific case of M1 stimulation, this is now well established. Here, we take this argument one step further, demonstrating that it can also lead to widespread distal modulations of cortico-subcortical loops when participants are engaged in a relevant cognitive task, and that such modulations do not require the participant to engage with the said task while receiving the stimulation itself. Specifically, our predicted changes to thalamo-cortical dynamics induced by anodal M1-tDCS (as discussed above), are consistent with, and expand, the now widely reported effects on M1 excitability[14] as well as more recently described changes to BOLD signal[15,44,45] and functional connectivity at rest[46–48]. In contrast, the effect of cerebellar tDCS on neural dynamics is much less understood. As discussed above, cathodal cb-tDCS increased thalamic afferent excitation over M1. In contrast, anodal stimulation led to increased self-inhibition in both M1 and thalamus. These findings demonstrate that tDCS is able to modulate cerebellar-brain inhibition (CBI) in a polarity specific manner, in agreement with previous electrophysiological reports[16], as well as a recent report of local increased activation in the dentate nuclei after cathodal cb-tDCS during simple finger tapping[49]. Furthermore, for the first time, we provide a window into the specific functional dynamics mediating these effects.

Interestingly, against our prediction, cathodal tDCS over M1 also led to an increase in thalamic activation and in excitation from thalamus to M1, as compared to sham. These changes further support the already described complex effects that characterise this polarity [7]. Specifically, cathodal tDCS is known to produce more inconsistent behavioural results than anodal stimulation, although these inconsistencies are more common in cognitive than motor studies [50]. Interestingly, our cathodal M1-tDCS also increased the modulatory effect of the task over M1 (i.e., led to greater M1 inhibition during the move blocks), but this was not accompanied by reductions in motor performance in the task. We believe this suggests that the thalamic changes reflect a compensatory mechanism to overcome cortical inhibition caused by cathodal M1-tDCS and to maintain an acceptable level of motor performance. This is in line with earlier animal models suggesting sustained effects of tDCS that are characterised by the system trying to compensate and normalise its activity to baseline levels (see [51] as discussed in [52]). Similarly, in Experiment 2, cathodal stimulation over the cerebellum led to the expected increases in excitatory output from thalamus to M1, but also an unexpected increase in M1 self-inhibition. Once again, tDCS did not alter behavioural performance and thus we believe this cortical reduction also compensated for the excess excitation coming from the thalamus. Alongside determining whether these changes have a therapeutic effect, neuroimaging studies of tDCS in PDOC will help elucidate whether the effects of cathodal M1-tDCS and anodal cb-tDCS are indeed compensatory or can alter behaviour when a motor deficit is present. In either case, in showing polarity specific modulations for some but not all our results, our study speaks for the complexity of the effects of tDCS [39] and suggests that other active control conditions alongside polarity should be included in future studies.

Several limitations need to be acknowledged. First, the distribution of the current generated by conventional tDCS is characterised by very low spatial accuracy and can reach a widespread area beyond the intended target. As seen in the simulations provided in Fig. 1, our montages are no exception to this. Our simulations suggest that the delivered current did not reach the thalamus with either montage. Therefore, our reported effects for this structure are likely to be explained by modulations of network connectivity. However, simulations suggest that M1-tDCS generated similar levels of current in SMA to that of M1 itself, and thus we cannot rule out that some of our effects are mediated by SMA. In contrast, our modelled current distribution for cb-tDCS extended beyond cerebellum into occipital and ventral temporal regions. These areas are not associated with our motor command-following task and are therefore not likely to have driven our effects. In either case, while the lack of spatial specificity does not limit the potential clinical application of tDCS in PDOC, it should be considered when making inferences about causal links between elicited effects and specific brain areas. Future studies should consider using a montage targeting non motor regions to make stronger causal inferences about the role of specific areas. Additionally, high-definition tDCS (HD-tDCS) can achieve higher spatial precision [53]. However, as we have previously argued [30], the increased spatial precision of this method requires careful consideration of individual brain structure and tissue properties, especially in patients with severe brain damage, which might limit clinical applications of HD-tDCS in PDOC. Second, the effects of tDCS are highly dependent on the state of the target brain networks during stimulation [54], and are more effective when paired with a relevant task [55]. Using a task during stimulation also partially overcomes the above limitations in spatial accuracy in ensuring that the effects are maximal for the intended areas (amongst all areas receiving current). Additionally, while we encouraged our participants to remain awake and monitored them during the 20 minutes of tDCS, the lack of behavioural outputs inherent to rest scans precluded us from verifying their wakefulness levels. It is thus possible that some of our participants experienced variable levels of wakefulness that could result in further individual differences in their brain states. However, as discussed above, PDOC patients are unable to voluntarily engage in behavioural tasks and delivering the stimulation at rest remains the most feasible option. Future studies should consider alternative ways to modulate brain states when designing tDCS interventions for this challenging patient group (e.g., see [56]). Third, in Experiment 2, we increased our FOV to ensure a full coverage of the cerebellum for all participants, and this required a longer TR. The resulting reduced temporal resolution that resulted may have affected our sensitivity to detect BOLD changes, compared to Experiment 1 [57]. We note that when all trials were included (e.g., see Fig. S1) the activation patterns were similar across both experiments, but this difference in sensitivity should be considered when making comparative arguments about effectiveness across our two montages. Importantly, DCM provides a more complete and sensitive account of differences in regional activation and their interactions, and can thus more reliably detect group differences [58]. Future studies with larger cohorts are required to clarify whether our proposed montages can elicit robust changes at the GLM level also.

## 5. Conclusions

In summary, our results indicate that tDCS can successfully modulate long-range thalamo-cortical dynamics underlying behavioural responsiveness during command following. It is yet to be tested whether these effects can be replicated in PDOC patients themselves and whether this will result in measurable clinical effects. However, our methodology can be directly applied to investigate this, and in doing so, it opens new avenues to explore the mechanisms of tDCS interventions in this challenging population.

## Data availability statement

Processed data is available from the authors upon reasonable request. Please contact with any questions or requests.

## Declarations of interest

None

## Acknowledgements

This work was supported by generous funding from the Medical Research Council (MR/P02596X/1; DF-E). DA was supported by a scholarship from The Wellington Hospital and the University of Birmingham. We thank Prof Michael Stevens for their advice on data cleaning, and all the volunteers for their time.

## Author Contributions

DF-E designed the study and obtained funding. RJ, PT, and DFE collected the data. DA and DF-E analysed the data, interpreted the results, prepared the figures, and wrote the manuscript. CRM, PT contributed to the editing of the final draft. All authors approved the content of the manuscript.

## Supplementary Material

**Table S1.**
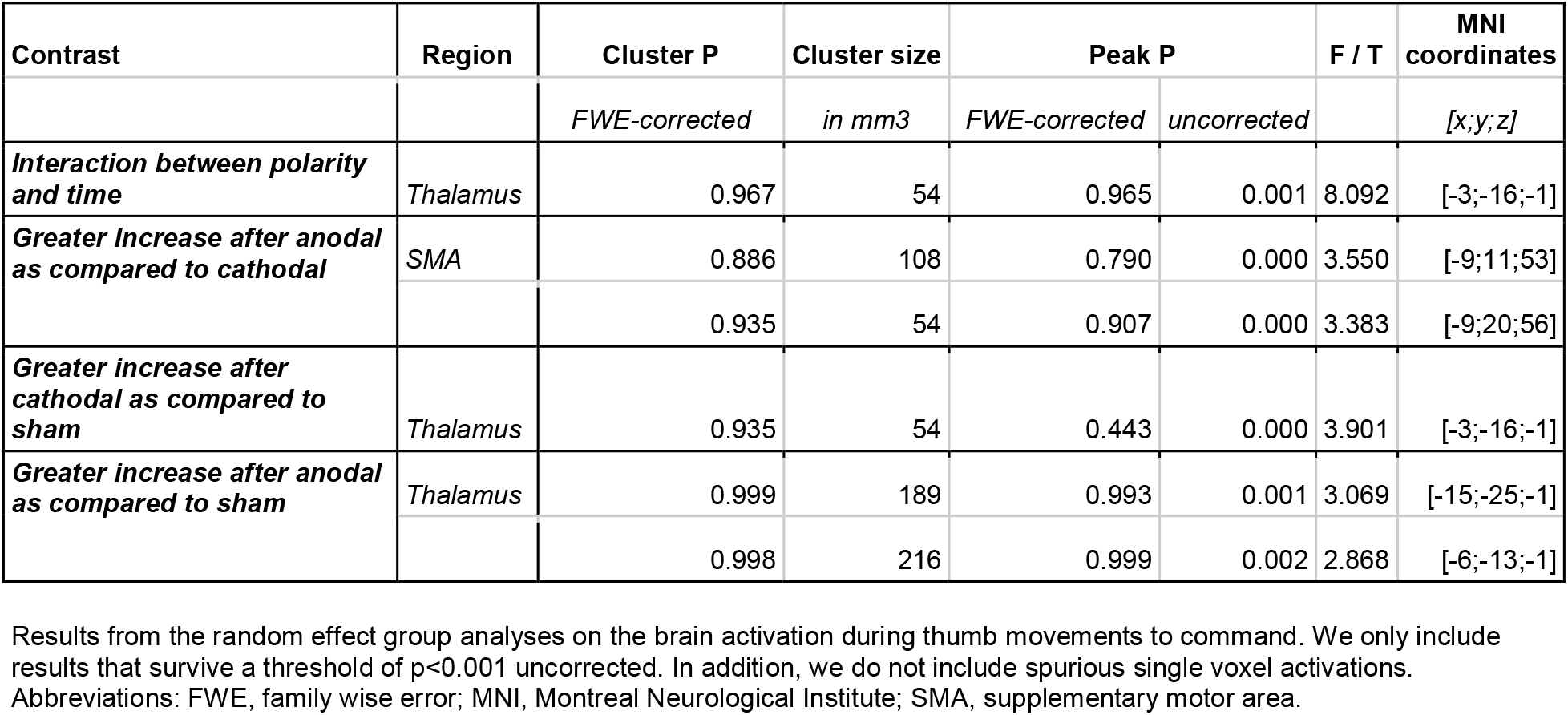
Effect of M1-tDCS on brain activation.

| Contrast | Region | Cluster P | Cluster size | Peak P |  | F / T | MNI coordinates |
| --- | --- | --- | --- | --- | --- | --- | --- |
|  |  | *FWE-corrected* | *in mm3* | *FWE-corrected* | *uncorrected* |  | *[x;y;z]* |
| **Interaction between polarity and time** | Thalamus | 0.967 | 54 | 0.965 | 0.001 | 8.092 | [-3;-16;-1] |
| **Greater Increase after anodal as compared to cathodal** | SMA | 0.886 | 108 | 0.790 | 0.000 | 3.550 | [-9;11;53] |
|  |  | 0.935 | 54 | 0.907 | 0.000 | 3.383 | [-9;20;56] |
| **Greater increase after cathodal as compared to sham** | Thalamus | 0.935 | 54 | 0.443 | 0.000 | 3.901 | [-3;-16;-1] |
| **Greater increase after anodal as compared to sham** | Thalamus | 0.999 | 189 | 0.993 | 0.001 | 3.069 | [-15;-25;-1] |
|  |  | 0.998 | 216 | 0.999 | 0.002 | 2.868 | [-6;-13;-1] |

Results from the random effect group analyses on the brain activation during thumb movements to command. We only include results that survive a threshold of $p < 0.001$ uncorrected. In addition, we do not include spurious single voxel activations. Abbreviations: FWE, family wise error; MNI, Montreal Neurological Institute; SMA, supplementary motor area.

**Table S2.** Effects of tDCS on behavioural metrics.

| Metrics | Polarity | Baseline | Post-tDCS | <u>t</u><sub>(baseline vs post-tDCS)</sub> | <u>p</u>-Holm<br><sub>(baseline vs post-tDCS)</sub> | <u>F</u><sub>(main effect time)</sub> | <u>p</u><sub>(main effect time)</sub> | <u>F</u><sub>(interaction)</sub> | <u>p</u><sub>(interaction)</sub> |
| --- | --- | --- | --- | --- | --- | --- | --- | --- | --- |
| Experiment 1 - M1-tDCS |  |  |  |  |  |  |  |  |  |
| Reaction time (<u>s</u>) | Anodal | 0.29 (± 0.06) | 0.27 (± 0.05) | <u>2.389</u> | <u>0.314</u> | <u>3.990</u> | <u>0.063</u> | 1.312 | 0.283 |
|  | Cathodal | 0.27 (± 0.05) | 0.26 (± 0.04) | <u>0.626</u> | <u>1.0</u> |  |  |  |  |
|  | Sham | 0.28 (± 0.05) | 0.28 (± 0.07) | <u>0.102</u> | <u>1.0</u> |  |  |  |  |
| Mean Velocity (<u>cm/s</u>) | Anodal | 8.39 (± 3.98) | 7.59 (± 2.81) | <u>2.697</u> | <u>0.145</u> | <u>04.430</u> | <u>0.051</u> | 1.939 | 0.160 |
|  | Cathodal | 7.71 (± 4.49) | 7.26 (± 3.83) | <u>0.868</u> | <u>1.0</u> |  |  |  |  |
|  | Sham | 6.96 (± 3.14) | 6.99 (± 2.83) | <u>0.120</u> | <u>1.0</u> |  |  |  |  |
| Peak acceleration (<u>m/s<sup>2</sup></u>) | Anodal | 31.05 (± 41.64) | 37.31 (± 56.13) | <u>0.312</u> | <u>1.0</u> | <u>0.616</u> | <u>0.444</u> | 0.636 | 0.536 |
|  | Cathodal | 32.81 (± 38.32) | 24.11(± 29.34) | <u>0.325</u> | <u>1.0</u> |  |  |  |  |
|  | Sham | 77.82 (± 177.01) | 111.39 (± 244.91) | <u>1.299</u> | <u>1.0</u> |  |  |  |  |
| Experiment 2 - cb-tDCS |  |  |  |  |  |  |  |  |  |
| Reaction time (<u>s</u>) | Anodal | 0.30 (± 0.04) | 0.28 (± 0.04 ) | 3.669 | 0.008** | <u>21.094</u> | <u><0.001**</u> | 0.951 | 0.395 |
|  | Cathodal | 0.29 (±0.03) | 0.28 (± 0.04) | <u>1.998</u> | <u>0.601</u> |  |  |  |  |
|  | Sham | 0.30 (± 0.05) | 0.29 (±0.05) | <u>1.885</u> | <u>0.601</u> |  |  |  |  |
| Mean Velocity<br>(*cm/s*) | Anodal | 7.44 (± 3.53) | 7.78 (± 3.98) | <u>0.626</u> | <u>1.0</u> | <u>0.286</u> | <u>0.598</u> | <u>1.106</u> | <u>0.340</u> |
|  | Cathodal | 7.11 (± 3.18) | 6.38 (± 2.77) | <u>-1.486</u> | <u>1.0</u> |  |  |  |  |
|  | Sham | 8.09 (± 2.73) | 8.09 (± 2.55) | <u>-0.007</u> | <u>1.0</u> |  |  |  |  |
| Peak acceleration<br>(*m/s<sup>2</sup>*) | Anodal | 39.29 (± 31.94) | 39.29 (± 33.62) | <u>0.001</u> | <u>1.0</u> | <u>1.823</u> | <u>0.191</u> | 0.404 | 0.670 |
|  | Cathodal | 51.19 (± 96.90) | 31.74 (± 15.27) | <u>-0.372</u> | <u>1.0</u> |  |  |  |  |
|  | Sham | 77.82 (± 141.91) | 60.96 (± 108.44) | <u>-1.046</u> | <u>1.0</u> |  |  |  |  |

Statistics for the post hoc (baseline vs post-tDCS) tests, main effect of time, and interaction between polarity (anodal, cathodal, sham) and time (baseline vs post-tDCS) on average reaction time, mean velocity and peak acceleration for Experiment 1 and Experiment 2. p-Holm value adjusted for comparing a family of 15; $p_{(\text{interaction})}$ uncorrected; \*\* $p < .01$ . Abbreviations: ms, milliseconds; cm, centimetres; s, seconds; m, metres.

**Table S3.** Blinding.

| Experiment | Active Stimulation | Sham | $\chi^2$ | p |
| --- | --- | --- | --- | --- |
| Experiment 1 | 31/43 | 15/22 | 0.9682 | 0.701 |
| Experiment 2 | 33/40 | 13/22 | 0.0869 | 0.263 |

Number of times that each type of stimulation was perceived as real, and statistics for the corresponding McNemar's Test. Active stimulation includes anodal and cathodal sessions.

**Figure S1.**
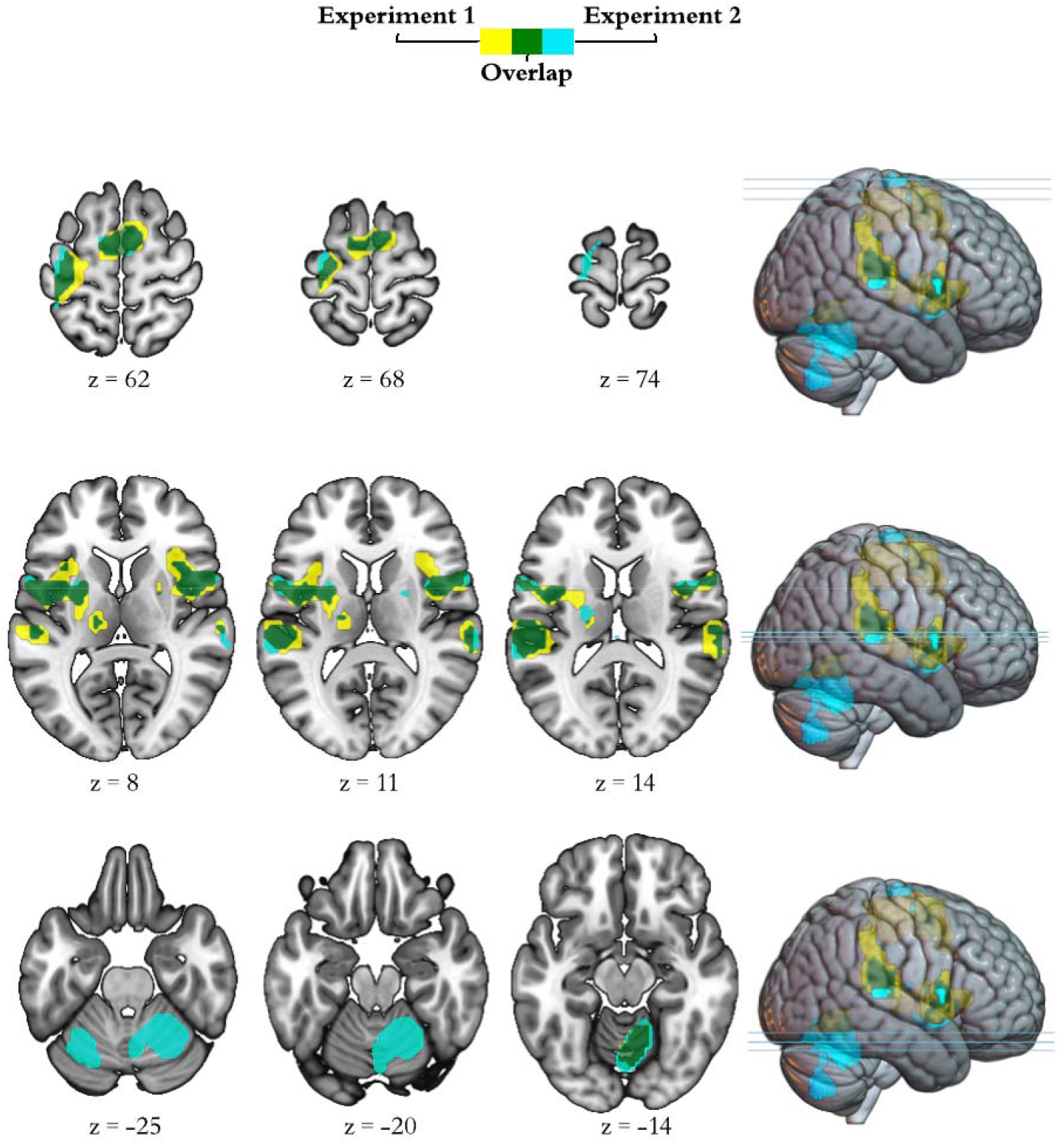
Brain activation during command following across trials. The insets display group general linear model differences between ‘move’ and ‘rest’ blocks in Experiment 1 (yellow) and Experiment 2 (light blue), across all trials included in the ANOVA (positive effect of task). The overlap across experiments appears in green. For display purposes, activation maps are shown at a FWE p<0.05 and rendered on a standard template (152 template in MRIcroGL). We display whole brain results as per request during peer review. z indicates the Montreal Neurological Institute z coordinate.

## Notes

### Competing Interest Statement

The authors have declared no competing interest.

### Summary of Updates

GLM methods and results

